# Studying Human Behaviour with Virtual Reality: The Unity Experiment Framework

**DOI:** 10.1101/459339

**Authors:** Jack Brookes, Matthew Warburton, Mshari Alghadier, Mark Mon-Williams, Faisal Mushtaq

## Abstract

Virtual Reality systems offer a powerful tool for human behaviour research. The ability to create three-dimensional visual scenes and measure responses to the visual stimuli enables the behavioural researcher to test hypotheses in a manner and scale that were previously unfeasible. For example, a researcher wanting to understand interceptive timing behaviour might wish to violate Newtonian mechanics, so objects move in novel 3D trajectories. The same researcher may wish to collect such data with hundreds of participants outside the laboratory, and the use of a VR headset makes this a realistic proposition. The difficulty facing the researcher is that sophisticated 3D graphics engines (e.g. Unity) have been created for game designers rather than behavioural scientists. In order to overcome this barrier, we have created a set of tools and programming syntaxes that allow logical encoding of the common experimental features required by the behavioural scientist. The Unity Experiment Framework (UXF) allows the researcher to readily implement several forms of data collection, and provides researchers with the ability to easily modify independent variables. UXF does not offer any stimulus presentation features, so the full power of the Unity game engine can be exploited. We use a case study experiment, measuring postural sway in response to an oscillating virtual room, to show how UXF can replicate and advance upon behavioural research paradigms. We show that UXF can simplify and speed up development of VR experiments created in commercial gaming software and facilitate the efficient acquisition of large quantities of behavioural research data.

## Introduction

Virtual Reality (VR) systems are opening up new opportunities for behavioural research as they allow visual (and auditory) stimuli to be displayed in 3D computer generated environments that can correspond to the participant’s normal external Cartesian space, but which do not need to adhere to the rules of Newtonian mechanics (Wann & Mon-Williams, 1996). Moreover, VR systems support naturalistic interactions with virtual objects and can provide precise measures of the kinematics of the movements made by adults and children in response to displayed visual stimuli. In addition, the relatively low cost and portability of these systems lowers the barriers to performing research in non-laboratory settings.

The potential advantages of VR in behavioural research have been recognised for at least two decades (e.g. Loomis, Blascovich, & Beall, 1999) but recent advantages in technology and availability of hardware and software are making VR a feasible tool for all behavioural researchers (rather than a limited number of specialist VR labs). For example, researchers can now access powerful software engines that allow the creation of rich 3D environments. One such popular software engine is Unity (alternatively called Unity3D; Unity Technologies, 2018). Unity is a widely used 3D game engine for developing video games, animations and other 3D applications and it is growing in its ubiquity. It is increasingly being used in research settings as a powerful way of creating 3D environments for a range of applications (e.g. psychology experiments, surgical simulation, rehabilitation systems). The recent popularity of VR head-mounted displays has meant that Unity has become widely used by games developers for the purpose of crating commercial VR content. Unity has well developed systems in place for rich graphics, realistic physics simulation, particles, animations and more. Nevertheless, it does not contain any features specifically designed for the needs of human behaviour researchers. We set out to produce an open source software resource that would empower researchers to exploit the power of Unity for behavioural studies.

A literature search of human behavioural experiments reveals that experiments are often defined by a common model, one that more easily allows researchers to exercise the scientific method. Experiments are often composed of trials, where trials can be defined as an instance of a scenario. Trials are usually composed of a stimulus and a human response and are a basic unit of behavioural experiments. Trials can be repeated many times for a single participant, increasing the signal-to-noise ratio of measurements, or allowing the study of human behaviour over time (e.g. adaptation and learning). Blocks can be defined as a grouping of trials that share something in common; comparing measures between blocks allows the examination of how substantial changes to the scenario affect the response. A session is a single iteration of the task with a participant. Defining an experiment in such a session-block-trial model (**Figure 1**) allows the definition and communication of an experimental design without ambiguity.

**Figure 1.**
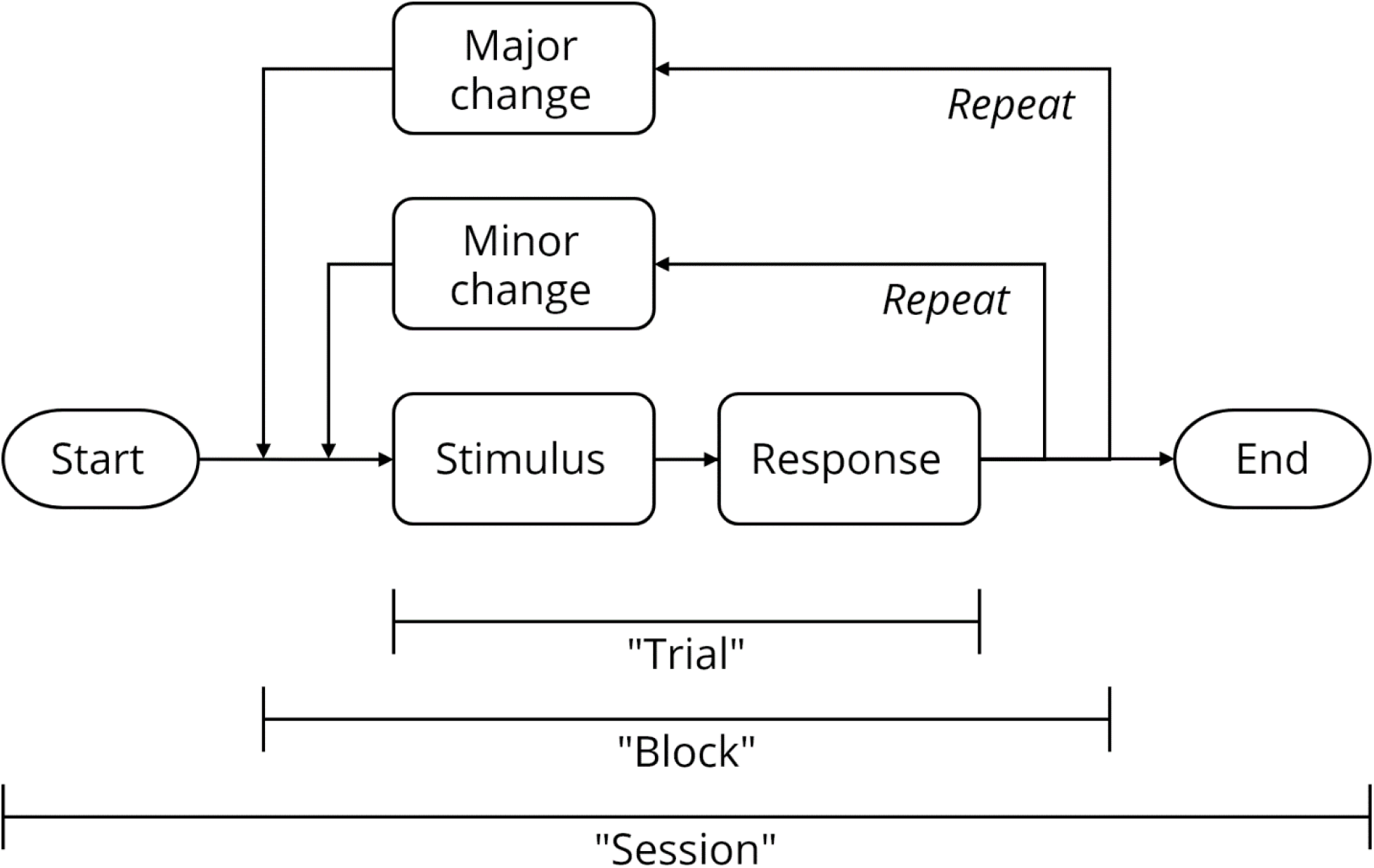
Structure of typical human behaviour experiments, in the session-block-trial model. Many experiments comprise multiple repetitions of trials. Between trials, only minor changes are made. A substantial change of content in the trial is often described as creating a new “block”. A single iteration of a task by a participant is called a session.

The use of this session-block-trial model in computer-based experiments affords a certain type of system design structure that mirrors the model itself. Typically, the code produced for an experimental task consists of a loop, where the process of presenting a stimulus and measuring a response is repeated many times, sometimes changing the parameters between loop iterations. The popularity of this experimental architecture means that researchers have attempted to provide tools that allow the development of tasks without the need to ‘reinvent the wheel’. Relatedly, development of the stimuli for software experiments is often difficult without knowledge of low-level computer processes and hardware. Thus, several software packages have been released which aim to make the stimuli themselves easier to specify in code. There is some crossover between these two types of packages, some focus only on stimuli whilst others also provide high-level ways to define the trials and blocks of the experiment and we briefly consider some of the most commonly used tools next.

PsychToolbox (Brainard, 1997) is a software package for MATLAB that allows researchers to program stimuli for vision experiments, providing the capability to perform low-level graphics operations but retaining the simplicity of the high-level interpreted MATLAB language. PsychoPy (Peirce, 2007) is an experimental control system that provides a means of using the Python programming language to systematically display stimuli to a user with precise timing. It consists of a set of common stimulus types, built-in functions for collection and storage of user responses/behaviour, and means of implementing various experimental design techniques (such as parameter staircases). PsychoPy also attempts to make research accessible for non-programmers with its ‘builder’, a GUI (graphical user interface) that allows development of experiments with little to no computer programming requirements.

The graphics processes for immersive technologies are significantly more complex than those required for two dimensional displays. In VR, it is difficult to think of stimuli in terms of a series of coloured pixels. The additional complexity includes a need for stimuli to be displayed in apparent 3D to simulate the naturalistic way objects appear to scale, move and warp according to head position. Unity and other game engines have the capacity to implement the complex render pipeline that can accurately display stimuli in a virtual environment; current academic focused visual display projects may not have the resources to keep up with the evolving demands of immersive technology software. Vizard (WorldViz, 2018), Unreal Engine (Epic Games, 2018), and open-source 3D, game engines such as Godot (Godot, 2018) and Xenko (Xenko, 2018) are also feasible alternatives to Unity, but Unity may still be a primary choice for researchers because of its ease of use, maturity, and widespread popularity.

### The Unity Experiment Framework (UXF)

To provide behavioural researchers with the power of Unity and the convenience of programmes such as PsychoPy, we created the Unity Experiment Framework (UXF). UXF is a software framework for the development of human behaviour experiments with Unity and the main programming language it uses, C#. UXF takes common programming concepts and features that are widely used, and often re-implemented for each experiment, and implements them in a generic fashion (**Table 1**). This gives researchers the tools to create their experimental software without the need to re-develop this common set of features. UXF aims to specifically solve this problem, and overtly excludes any kind of stimulus presentation system, with the view that Unity (and its large asset developing community) provides all the necessary means to implement any kind of stimulus or interaction system for an experiment. In summary, UXF provides the ‘nuts and bolts’ that work behind the scenes of an experiment developed within Unity.

**Table 1.** Common experiment concepts and features which are represented in UXF

| Concept | Description |
| --- | --- |
| Trial | The base unit of experiments. A trial is usually a singular attempt at a task by a participant after/during the presentation of a stimulus. |
| Block | A set of trials – often used to group consecutive trials that share something in common. |
| Session | A session encapsulates a full “run” of the experiment. Sessions are usually separated by a significant amount of time and could be within subjects (for collection of data from a singular participant over several sessions) and/or between subjects (for collection of data from several participants each carrying out a single session). |
| Settings | Settings are parameters or variables for an experiment, block, or trial, usually predetermined, that quantitatively define the experiment. Settings are useful for defining the experimental manipulation (i.e. the independent variables). |
| Behavioural data | We perform an experiment to measure the effect of an independent variable on a dependent variable. Behavioural data collection allows for the collection of measured values of dependent variables on a trial-by-trial basis. For example, we may wish to collect the response to a multiple-choice question, or the distance a user throws a virtual ball. |
| Continuous data | Within a trial, we may want to measure a value of one or more parameters over time. Most commonly we want to record the position and rotation of an object within each trial. This could be an object that is mapped to a real-world object (e.g. participant head, hands) or a fully virtual object (virtual ball in a throwing experiment). Position and rotation of an object is the main use case but UXF supports measurement of any parameter over time (e.g. pressure applied to a pressure pad). |
| Participant information | There may be other variables that we cannot control within the software which we may wish to measure to record to examine its relationship to the result. For example, age or gender of the participant. |

### Experiment structure

UXF provides a set of high-level objects that directly map onto how we describe experiments. The goal is to make the experiment code more readable and avoid the temptation for inelegant if-else statements in the code as the complexity increases. Session, blocks, trials are our ‘objects’ which can be represented within our code. The creation of a session, block or trial automatically generates properties we would expect them to have – for example each block has a block number, each trial has a trial number. These numbers are automatically generated as positive integers based on the order in which they were created. Trials contain functionality such as ‘begin’ and ‘end’ which will perform useful tasks implicitly in the background, such as recording the timestamp when the trial began or ended. Trials and blocks can be created programmatically, meaning UXF can support for any type of experiment structure, including staircase or adaptive procedures.

### Measuring dependent variables

While the trial is ongoing, at any point we can add any observations to the results of the trial, which will be added to the behavioural data .CSV output file at the end of the session. Additionally, we can continuously log a variable over time at the same rate as the display refresh frequency (90Hz in most currently-available commercial VR HMDs). The main use case of this is where the position and rotation of any object in Unity can be automatically recorded on a per-trial basis, saving a single .CSV file for each trial of the session. This allows for easy cross-referencing with behavioural data. All data files (behavioural, and continuous) are stored in a directory structure organised by *experiment > participant > session number*.

### Setting independent variables

Settings can be used to attach values of an independent variable to an experiment, session, block, or trial. Settings have a cascading effect, whereby one can apply a setting to the whole session, a block or a single trial. When attempting to access a setting, if it has not been assigned in the trial, it will attempt to access the setting in the block. If it has not been assigned in the block, it will search in the session (Figure 2). This allows users to very easily implement features common to experiments, such as “10% of trials contain a different stimulus”. In this case, one could assign a “stimulus” setting for the whole session, but then assign 10% of the trials with a different value for a “stimulus” setting.

**Figure 2.**
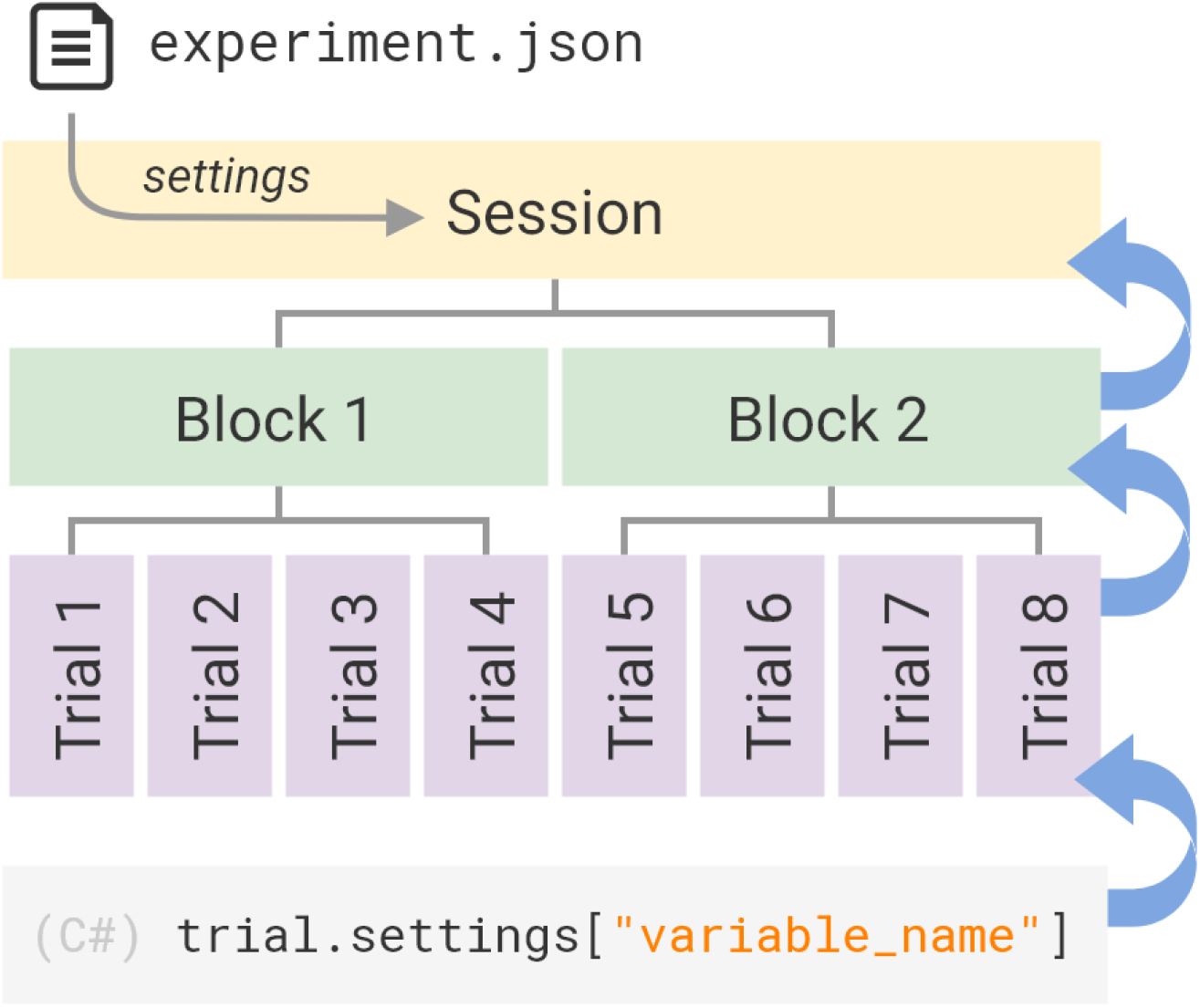
The UXF Settings system. Independent variables that we change in order to iterate a design of an experiment, or to specify the experimental manipulation itself, can be written in a human-readable .json file. Settings can also be programmatically accessed or created at trial, block or session level. Where a setting has not been specified, the request cascades up and searches in the next level above. This allows both “gross” (e.g. to a whole session) or “fine” (e.g. to a single trial) storage of parameters within the same system.

Settings are also a useful feature for allowing for changing experimental parameters without modifying the source code. A simple text file (.JSON format) can be placed in the experiment directory which will be read upon the start of a session, and its settings applied to that session. This system speeds up the iteration time during the process of designing the experiment; the experimenter can change settings from this file and see their immediate effect without changing any of the code itself. It also allows multiple versions of the same experiment (e.g. different experimental manipulations) to be maintained within a single codebase using multiple settings files. One of these settings profiles can be selected by the experimenter on launching the experiment task.

### Experimenter User Interface

UXF includes an (optional) experimenter user interface (UI) to allow selection of a settings profile, and inputting additional participant information, such as demographics. Information the experimenter wishes to collect is fully customisable. The UI includes support for a “participant list” system, whereby participant demographic information is stored in its own CSV file. As new participants perform the experiment, their demographic information is stored in the list. This allows participant information to be more easily shared between sessions or even separate experiments – instead of having to input the information each time, the experimenter can select any existing participant found in the participant list via a drop-down menu.

**Figure 3.**
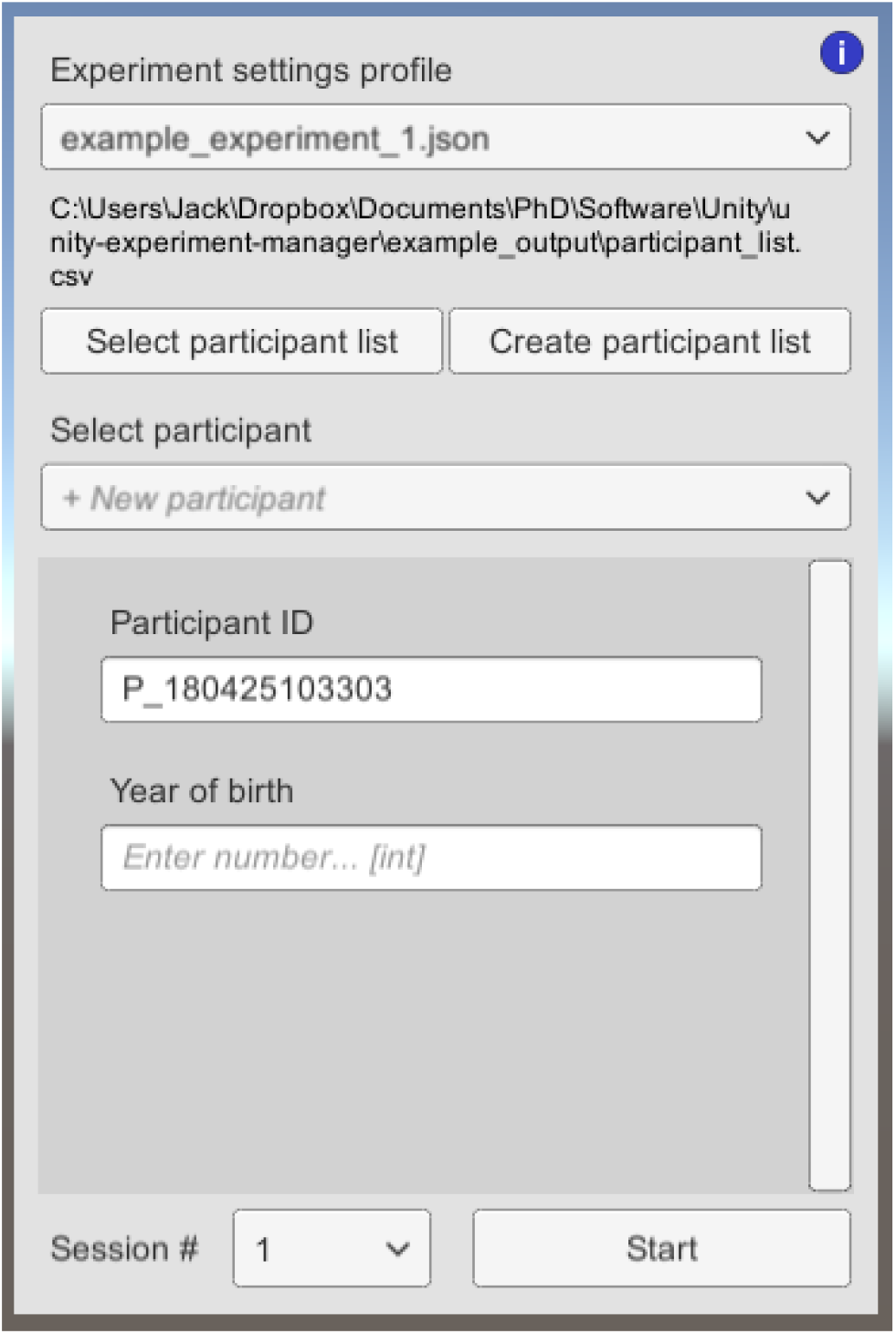
Screenshot of the experimenter user interface.

### Example

Below is an example of the C# code used to generate a simple 2 block, 10 trial experiment where the participant is presented with a number *x* and they must input the doubled value (2*x*).

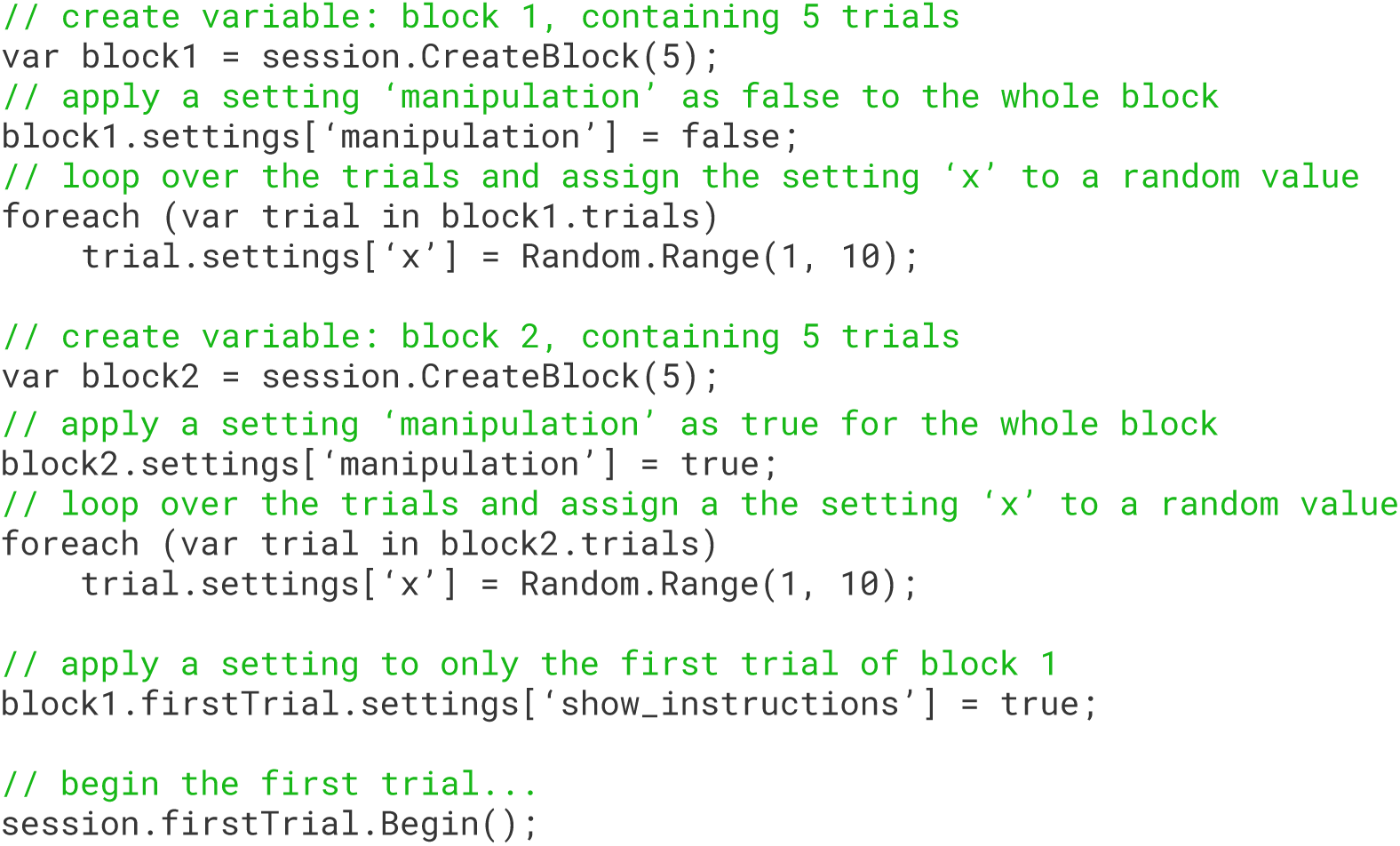

Elsewhere in our project, we must define what happens when we begin the trial (such as making the value of *x* appear for the participant), and mechanisms to retrieve the participant’s response for the trial (participant’s calculated value of 2*x*). These are to be created with standard Unity features for making objects appear in the scene, collecting user response via keyboard input, etc. The resulting behavioural data .CSV file would be automatically generated and saved (**Table 2**). A typical structure of a task developed with UXF is shown in **Figure 4**.

**Table 2.** Example behavioural data output. Columns not shown include participant ID, session number, and experiment name.

| trial_num | block_num | start_time | end_time | manipulation | x | response |
| --- | --- | --- | --- | --- | --- | --- |
| 1 | 1 | 0.000 | 1.153 | FALSE | 8 | 16 |
| 2 | 1 | 1.153 | 2.112 | FALSE | 3 | 6 |
| 3 | 1 | 2.112 | 2.950 | FALSE | 4 | 8 |
| 4 | 1 | 2.950 | 3.921 | FALSE | 7 | 14 |
| 5 | 1 | 3.921 | 4.727 | FALSE | 4 | 8 |
| 6 | 2 | 4.727 | 5.826 | TRUE | 9 | 18 |
| 7 | 2 | 5.826 | 6.863 | TRUE | 5 | 10 |
| 8 | 2 | 6.863 | 7.693 | TRUE | 10 | 20 |
| 9 | 2 | 7.693 | 8.839 | TRUE | 6 | 12 |
| 10 | 2 | 8.839 | 9.992 | TRUE | 3 | 6 |

**Figure 4.**
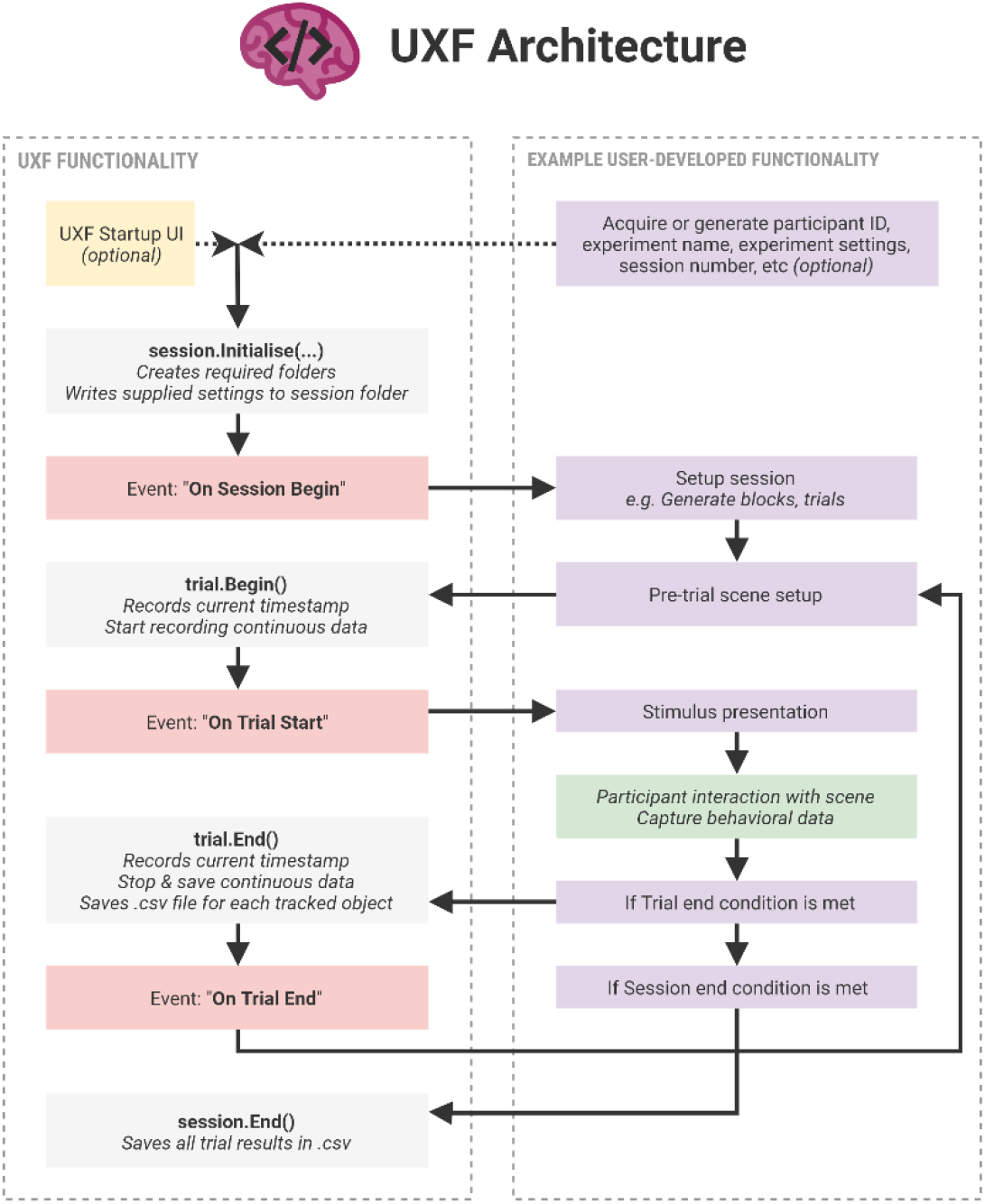
Structure of a typical task developed with UXF. The left panel shows functionality present in UXF, with functionality a researcher is expected to implement shown on the right panel. The framework features several “events” (shown in red) which are invoked at different stages during the experiment; these allow developers to easily add behaviours that occur at specific times, for example presenting a stimulus at the start of a trial.

### Multithreading file I/O

Continuous measurement of variables requires large amounts of data to be collected over the course of the experiment. When using a VR head-mounted display, it is essential to maintain a high frame rate and keep stutters to a minimum to minimise the risk of inducing sickness or discomfort on the participant. Handling of tasks such as reading and writing to file may take several milliseconds or more depending on operating system background work. Constant data collection (particularly when tracking the movement of many objects in the scene) and writing these data to file therefore poses a risk of dropping the frame rate below acceptable levels. The solution is to create a multi-threaded application which allows the virtual environment to continue to be updated whilst data are being written to files simultaneously in a separate thread. Designing a stable multithreaded application imparts additional technical requirements on the researcher. UXF abstracts file I/O away from the developer, performing these tasks automatically, with a multithreaded architecture working behind the scenes. Additionally, the architecture contains a queueing system, where UXF queues up all data tasks and writes the files one-by-one, even halting the closing of the program to finish emptying the queue if necessary.

### Cloud-based experiments

UXF is a standalone, generic project, and so it does not put any large design constraints on developers using it. This means that UXF does not have to be used in a traditional lab-based setting, with researchers interacting directly with participants; it can be used for data collection opportunities outside of the lab, by embedding experiments within games or apps that a user can partake in at their discretion. Data are then sent to a web server where it can later be downloaded and analysed by researchers (Figure 5). Recently these cloud-based experiments have become a viable method of performing experiments on a large scale.

**Figure 5.**
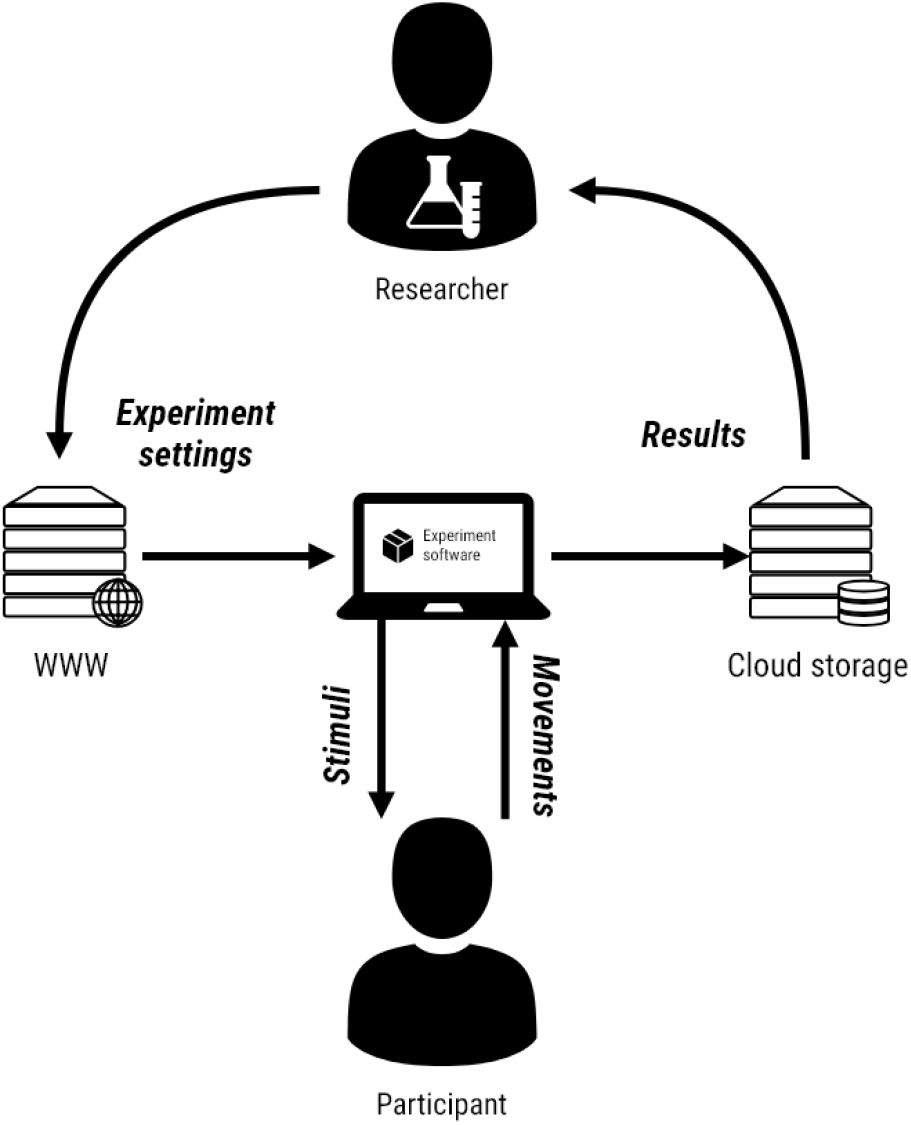
Experiment in the cloud. A piece of software developed with UXF can be deployed to an internet connected device. Researchers can modify experiment settings to test different experimental manipulations over time, which are downloaded from the web by the client device upon running a UXF experiment. As the participant partakes in the experiment, they are presented with stimuli, and their movements are recorded in the form of behaviours/responses or continuous measurement of parameters like hand position. Their results are automatically and securely streamed up to a server on the internet, of which the researcher can periodically retrieve data from.

UXF can be used in cloud-based experiments (**Figure 5**) using two independent pieces of software that accompany UXF:

1. *UXF S3 Uploader* allows all files that are saved by UXF (behavioural data, continuous data, logs) to be additionally uploaded to a location in Amazon’s Simple Storage Service as setup by a researcher. This utilizes existing UXF functionally of setting up actions for after a file has been written; and so a developer could potentially implement uploading the files to any other storage service.
2. *UXF Web Settings* replaces the default UXF functionality of selection of experiment settings via a user interface, to the settings being accessed automatically from a web URL by the software itself. This allows a deployed experiment (e.g. via an app store, or simply transferring an executable file), to be remotely altered by the researcher, without any modification to the source code. Settings files are stored in json format and would usually be of a very small file size so can be hosted online cheaply and easily.

A developer can implement neither, either, or both, depending on the needs of the research. For lab-based experiments, neither are required. For experiments without any need to modify settings afterwards, but with the requirement of securely backing up data in the cloud, (1) can be used. If a researcher wants to remotely modify settings but has physical access to the devices to retrieve data, (2) can be used. For a fully cloud-based experiment without direct researcher contact with the participant both (1) and (2) can be used. This has been successfully tried and tested in the context of a museum exhibition, where visitors could take part in VR experiments, with the recorded data being uploaded to the internet. Both UXF S3 Uploader and UXF Web Settings are available as open source Unity packages.

### Case study

One classic question in human behavioural research has related to the information used by adults and children when maintaining posture (Thomas & Whitney 1959; Edwards 1946). To investigate the contribution of kinaesthetic and vision information when both are available, four decades ago Lee and Aronson (1975) used a physical ‘swinging’ room to perturb the visual information provided by the walls and ceiling whilst leaving the kinaesthetic information unaffected (only the walls and ceiling swung and the floor did not move). This experiment demonstrated the influence of vision on posture but the scale of the apparatus meant that it could only ever be implemented in a laboratory setting. The approach was also subject to measurement errors and researcher bias (Wann, Mon-Williams & Rushton 1998). More recently, conventional computer displays have been used to explore the impact of vision on posture (e.g. Villard et al 2008) and this method has addressed issues of measurement error and researcher bias but still remains confined to the laboratory.

The ability to create a virtual swinging room in a VR environment provides a test case for the use of UXF in supporting behavioural research and provides a proof-of-concept demonstration of how large laboratory experiments can be placed within a non-laboratory setting. Here, we used the head tracking function as a proxy measure of postural stability (as decreased stability would be associated with more head sway; Flatters et al., 2014). In order to test the UXF software, we constructed a simple experiment with a within-participant component (whether the virtual room was stationary or oscillating) and a between-participant factor (adults vs children). We then deployed the experiment in a museum with a trained demonstrator and remotely collected data on one hundred participants.

The task was developed in the Unity game engine with UXF handling several aspects of the experiment including; Participant information collection, Settings, Behavioural data and Continuous data. *Participant information collection*: The UXF built-in user interface was used to collect a unique participant ID as well as the participant’s age and gender. This information was stored in a CSV participant list file. This list was subsequently updated with participant height and arm-span as they were collected in the task. *Settings*: A settings file accompanied the task that allowed modification of the assessment duration as well as the oscillation amplitude and period without modifying the code. Settings for each trial were used to construct the environment to facilitate the requested trial condition. *Behavioural data*: While there were no dependant variables that were directly measured on each trial, the UXF behavioural data collection system output a list of all trials that were run in that session, as well as the vision condition for that trial. *Continuous data*: UXF was configured to automatically log the HMD position over time within each trial, which was then used offline for the stability measure calculation. UXF split the files with one file per trial which was designed to make it easy to match each file with the trial condition the file was collected under.

## Methods

Fifty children (all under <16 years of age; mean age: 9.6 years; SD: 2.0 years) and 50 adults (mean age: 27.5 years; SD: 13.2 years) took part in the study. Participants were recruited from either the University of Leeds participant pool (adults), or were attendees at the Eureka! Science museum (children and adults) and provided full consent. A gaming-grade laptop (Intel Core i5-7300HQ, Nvidia GTX 1060) in addition to a VR HMD (Oculus Rift CV1) and the SteamVR API, a freely available package independent of UXF (Valve Corporation, 2018) were used to present stimuli and collect data. The HMD was first calibrated using the built-in procedure, which set the virtual floor level to match the physical floor.

After explaining task requirements, the demonstrator put the HMD on the participant’s head (over spectacles if necessary) and adjusted it until the participant reported it was comfortable and they could see clearly. Participants were then placed in the centre of a simple virtual room (height: 3m, width: 6m, depth: 6m) with textured walls and floors (Figure 6). Height was measured as vertical distance from the floor to the “centre eye” of the participant (as reported by the SteamVR API) and this value was used to place a fixation cross on the wall at the participant’s height.

**Figure 6.**
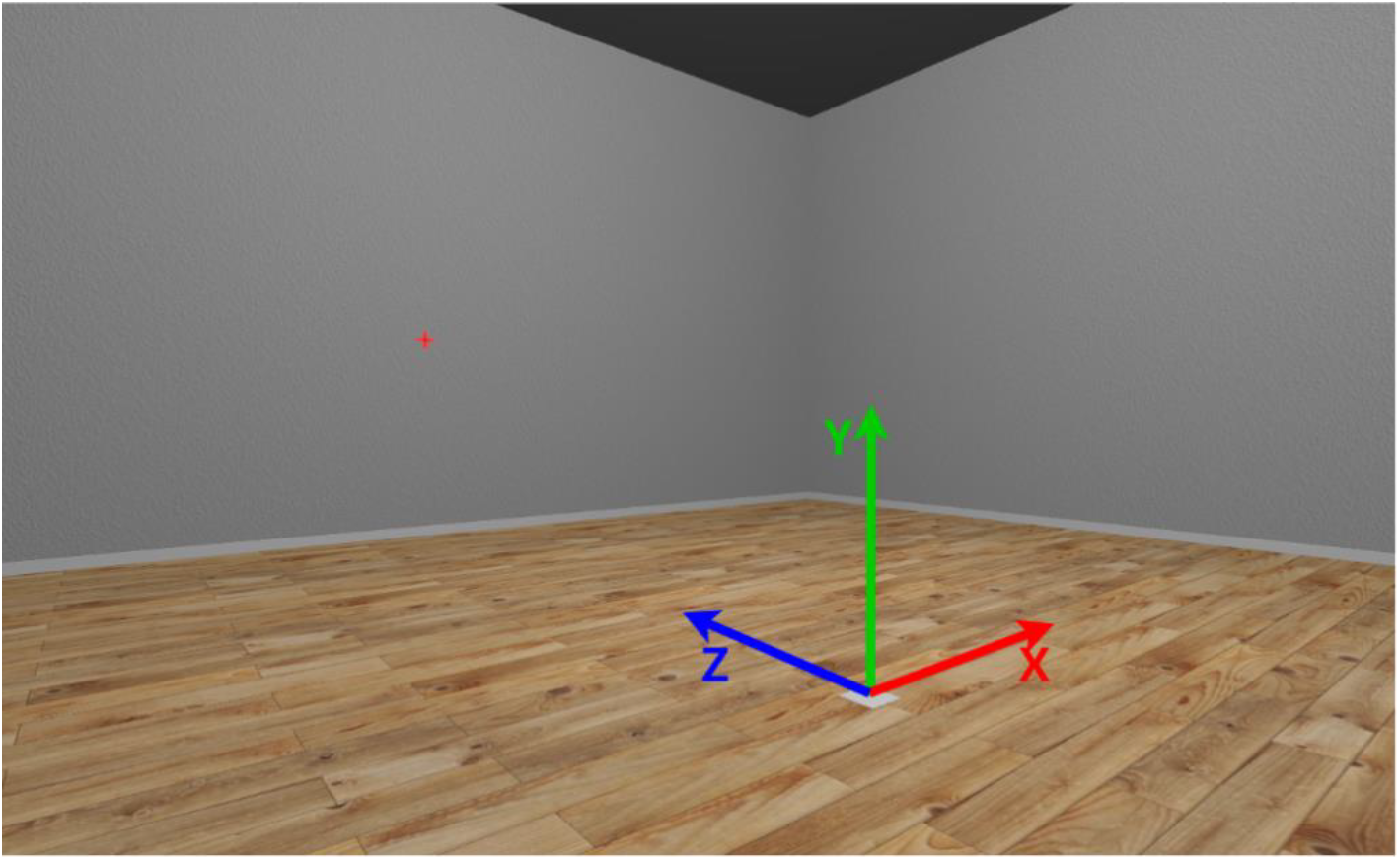
Screenshot from inside the virtual room. Arrows indicate the three axes as well as the origin. The red fixation cross is shown on the wall.

The task comprised two 10 second trials performed in a random order. The *normal* condition asked participants to stand still and look at a fixation cross placed on the wall. In the *oscillating* condition, the participants were given the same instructions, but the virtual room oscillated in a sinusoidal fashion (rotating around the x axis) with an amplitude of 5° and a frequency of 0.25Hz. The oscillation was performed about the point on the floor at the centre of the room, in effect keeping the participant’s feet fixed in-place. Participants were not explicitly informed about the room oscillation. The position of the HMD inside the virtual room was logged at a rate of 90Hz during each of the two trials. The path-length of the head was used as a proxy measure of postural stability (sum of all point-to-point distances over a trial).

## Results

No participants reported any feelings of sickness or discomfort during or after taking part in the task. A mixed-model design ANOVA (2 [Age: Adult vs Children] x 2 Vision Condition [Normal vs. Oscillating]) found no interaction, F(2, 98) = 0.34, p = .562, *η*^2^G = .001, but revealed main effects of Vision, F(2, 98) = 7.35, p = .008, *η*^2^G = .016 and Age, F(1, 98) = 9.26, p = .003, *η*^2^G = .068, thus replicating previous work on the contribution of visual information on postural stability (Flatters et al., 2014).

**Figure 7.**
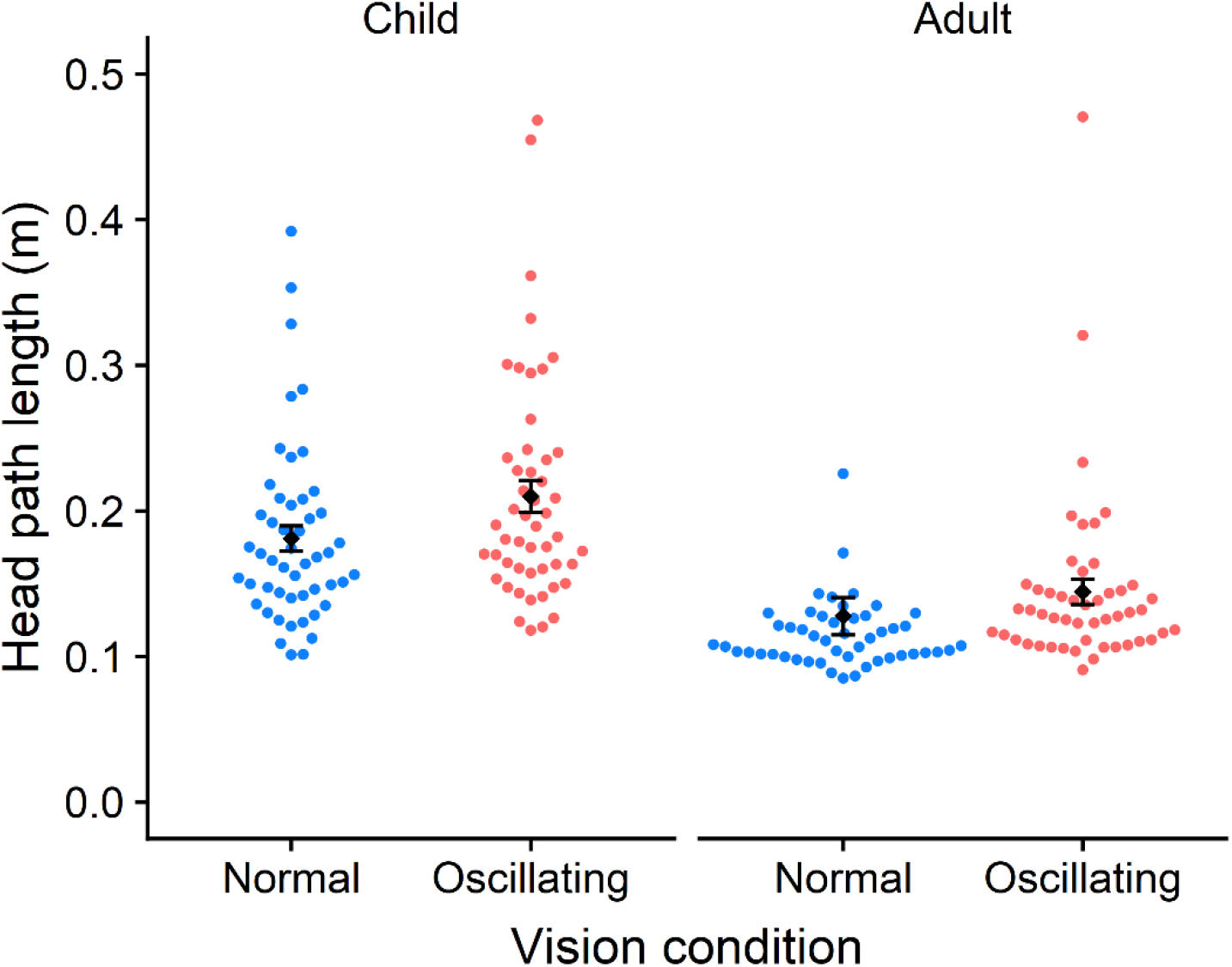
Head path length (higher values indicating worse postural stability) as a function of vision condition. The two conditions were ‘normal’ (static virtual room) and ‘oscillating’ (oscillating virtual room). Postural stability was indexed by the path length of head movement in meters (measured over a 10 second period). Adults showed significantly different path length overall compared to children (shorter – indicating greater stability). Error bars represent +/-1 SEM.

## Summary

We have created an open source resource that enables researchers to use the powerful games engine of Unity when designing experiments. We tested the usefulness of UXF by designing an experiment that could be deployed within a museum setting. We found that UXF simplified the development of the experiment and produced measures in the form of data files that were in a format that made subsequent data analysis straight forward. The data collected were consistent with the equivalent laboratory-based measures (reported over many decades of research) whereby children showed less postural stability than adults, and where both adults and children showed greater sway when the visual information was perturbed. There are likely to be differences in the postural responses of both adults and children within a virtual environment relative to a laboratory setting and we would not suggest that the data are quantitatively similar between these settings. Nonetheless, these data do show that remotely deployed VR systems can capture age differences and detect the outcomes of an experimental manipulation.

Our planned work includes maintaining the software for compatibility with future versions of Unity, and refactoring UXF so that it works on a wider range of platforms (e.g. mobile devices, web browsers, augmented reality devices, standalone VR headsets). Features may be added or modified if a clear need for a feature arises. The project is open source, thus allowing researchers in the field to implement and share such additions.

## Availability

UXF is freely available to download via GitHub as a Unity Package (github.com/immersivecognition/unity-experiment-framework), and currently can be integrated into Unity tasks built for Windows PCs. Documentation and support is available on the GitHub wiki (github.com/immersivecognition/unity-experiment-framework/wiki). The package is open sourced under the MIT licence. Related packages UXF S3 Uploader and UXF Web Settings are available via the same GitHub link.

## Acknowledgements

The authors would like to thank Almanzo McConkey and Andrea Loriedo for their feedback on earlier versions of the software. Authors F.M and M.M-W hold Fellowships from the Alan Turing Institute. M.W., F.M. and M.M-W are supported by a Research Grant from the EPSRC (EP/R031193/1).

